# Complete Chloroplast Genome of *Castanopsis sclerophylla* (Lindl.) Schott: Genome Structures, Comparative and Phylogenetic Analysis

**DOI:** 10.1101/540617

**Authors:** Xuemin Ye, Dongnan Hu, Yangping Guo, Rongxi Sun

## Abstract

*Castanopsis sclerophylla* (Lindl.) Schott is an important species of evergreen broad-leaved forest in subtropical area and has important ecological and economic value. However, there are little studies on its chloroplast genome. In this study, the complete chloroplast genome sequences of *C. sclerophylla* was reported based on the Illumina Hiseq 2500 platform. The complete chloroplast genome of *C. sclerophylla* was 160,497bp, including a pair of inverted repeated (IRs) regions (25,675bp) that were separated by a large single copy (LSC) region of 90,255bp, and a small single copy (SSC) region of 18,892bp. The overall GC content of chloroplast genome was 36.82%. A total of 131 genes were found, of these 111 genes were unique and annotated, including 79 protein-coding genes, 27 transfer RNA genes (tRNAs), and four ribosomal RNA genes (rRNAs). Twenty-one genes were found to be duplicated in the IR regions. Comparative analysis indicated that IR contraction might be the reason for the relatively smaller chloroplast genome size of *C. sclerophylla* compared with other three congeneric species. Sequence analysis detected that the LSC and SSC regions were more divergent than the IR regions within the *Castanopsis*, furthermore, a higher divergence was found in non-coding regions than in coding regions. The maximum likelihood (ML) phylogenetic analysis showed that these four species of the genus *Castanopsis* formed a monophyletic clade and that *C. sclerophylla* is closely related to *Castanopsis* hainanensis with strong bootstrap values. These results not only provide basic knowledge about characteristics of *C. sclerophylla* and also enhance our understanding of *Castanopsis* species evolution within the Fagaceae family. Meanwhile, these findings will contribute to the exploration, utilization and conservation genetics of *C. sclerophylla*.

## Introduction

*Castanopsis sclerophylla* (Lindl.) Schott. is a monoecious, and broad-leaved tree of the genus *Castanopsis* belongs to the Fagaceae family. The genus contains about 120 known species, of these species 58 are native to China, and 30 are endemic. However, this species is widely native to East and South Asia, and had been introduced to North America[1, 2]. In China, it is a canopy tree widely distributed in subtropical evergreen forests[3]. Its fruit and wood are valuable, and are regarded as landscape and ornamental tree because of glossy evergreen leaves and white flowers cover the tree [4]. In Jiangxi province, its fruit has been used to make special foods such as sheet jelly, bean curd, and bean vermicelli[5]. The previous researches mainly focused on natural regeneration[6], biomass[7], morphology[8], chemotaxonomical[9], genetic diversity[1, 10]. Currently, because of the tree’s economic value increasing, natural plants have been seriously destroyed by humans, and the number of natural trees is decreasing due to the slowly grow rate, this species is severely fragmented and constantly threatened, and need urgent conservation and restoration [11]. Phylogenetic evolution and population genomics knowledge are very vital to make effective conservation and management strategies. With the rapid development of next-generation sequencing technology, compared with the Sanger method, the chloroplast genome assembly has become cheaper and easier based on Illumina sequencing technology, comparative analysis of the complete chloroplast genome among closely related species has been proven to be a valid and most effective method for the study of evolutionary history, species conservation, and phylogenetic relationships [12–14].

Chloroplasts are essential organelles in plant cells which play very important roles for photosynthesis, carbon fixation, and pigments, starch, fatty acids, and amino acids synthesizing[15, 16]. Typically, the chloroplast genome of angiosperms is a highly conserved circular DNA ranging from 120 to 180 kb in higher plants, and has a typical quadripartite structure including a large single copy (LSC) region, a small single copy (SSC) region, and a pair of inverted repeats (IRs)[17]. The chloroplast genome encodes about 110 to 130 genes, including up to 80 unique protein-coding genes, four ribosomal RNAs (rRNAs), and approximately 30 transfer RNAs (tRNAs)[18]. In recent years, many complete chloroplast genome sequences of higher plants have been reported, which were used to study population structure and phylogenetic relationships.

In this study, we report the *C. sclerophylla* chloroplast genome using Illumina sequencing technology. This is the first comprehensive analysis of *C. sclerophylla* chloroplast genomes, in conjunction these whole chloroplast genome sequences of three congeneric species which were previously published. In addition, we also used 22 complete chloroplast genome sequences from GenBank to analyzed phylogenic relationships and infer the phylogenetic position of *C. sclerophylla*. These results not only provide basic knowledge about characteristics of *C. sclerophylla* and also enhance our understanding of *Castanopsis* species evolution within the Fagaceae family. Our data will contribute to understand the genetic resources and evolutionary based on the diversity in the chloroplast genome, and facilitate to the exploration, utilization and conservation genetics of *C. sclerophylla*.

## Materials and Methods

### Plant material, DNA Extraction and Sequencing

Fresh young leaves of *C. sclerophylla* were collected from the Jiangxi agricultural university arboretum in Nanchang, China (28°45’N, 115°49’E). Total genomic DNA was extracted using the Plant Genomic DNA Kit (TIANGEN, Beijing, China). Agarose gel electrophoresis and Microplate Spectrophotometer (Molecular Device, Sunnyvale, CA, USA) were used to measured DNA quality and concentration, respectively. Shotgun libraries with an average insert size of 350 bp were constructed using pure DNA and sequenced with an Illumina Hiseq 2500 platform. Approximately 10.44 GB of clean data from *C. sclerophylla* were yielded with 150 bp paired-end read lengths.

### Chloroplast Genome Assemblage and Annotation

The high throughput raw reads were trimmed by Fastqc. Next, trimmed paired-end reads and references (*C. hainanensis, C. echinocarpa*, and *C. concinna*) were used to extract chloroplast like reads. And then, these reads were assembled by NOVOPlasty[19]. NOVOPlasty assembled part reads and stretched as far as possible until a circular genome formed. Finally, high quality complete chloroplast genome was obtained. The assembled genome was annotated using CpGAVAS[20]. BLAST and Dual Organellar Genome Annotator (DOGMA) were used to check the annotation results [21]. These tRNAs were identified by the tRNAscanSE[22]. The circular gene maps of *C. sclerophylla* was drawn by the OGDRAWv1.2 program[23]. To analyze the variation of synonymous codon usage, MEGA7 was used to compute relative synonymous codon usage values (RSCU), codon usage, and the GC content[24].

### Comparative Analysis and Phylogenetic Analysis

MUMmer [25] was conducted for pairing sequence alignment of the chloroplast genomes. The mVISTA[26] program was used to compare the complete chloroplast genome of *C. sclerophylla* to other three published chloroplast genomes of the genus *Castanopsis*, i.e., *C. concinna* (NC_033409), *C. echinocarpa* (NC_023801), *C. hainanensis* (NC_037389) with the shuffle-LAGAN mode, adopting the annotation of *C. concinna* as a reference.

Phylogenies were constructed using the 19 cp genome of the Fagaceae species sequences from the NCBI Organelle Genome and Nucleotide Resources database: *Castanea_henryi* (NC_033881), *Castanea mollissima* (NC_014674), *Castanopsis concinna* (NC_033409), *Castanopsis echinocarpa* (NC_023801), *Castanopsis hainanensis* (NC_037389), *Fagus engleriana* (NC_036929), *Lithocarpus balansae*(NC_026577), *Quercus aliena*(NC_026790), *Quercus aquifolioides*(NC_026913), *Quercus baronii*(NC_029490), *Quercus dolicholepis*(NC_031357), *Quercus glauca*(NC_036930), *Quercus rubra*(NC_020152), *Quercus sichourensis*(NC_036941), *Quercus spinosa*(NC_026907), *Quercus tarokoensis*(NC_036370), *Quercus tungmaiensis*(NC_036936), *Quercus variabilis*(NC_031356), and *Trigonobalanus doichangensis*(NC_023959). The sequences were initially aligned using MAFFT[27]. Then, the visualization and manual adjustment of multiple sequence alignment was conducted in BioEdit[28]. The maximum likelihood (ML) analysis was conducted by RAxML web servers[29]. The bootstrap test was conducted 1000 times adopting Kimura 2-parameter model. *Corylus fargesii* (NC_031854), and *Eucalyptus umbra* (NC_022387) were used as the outgroups.

## Results and discussion

### Characteristics of C. sclerophylla cpDNA

A total of 65 million pair-end reads were obtained, with 10.44 Gb clean data. The complete chloroplast genome sequence of *C. sclerophylla* is 160,497 bp in length, which deposited in GenBank with accession number MK387847. Its genome possesses a typical quadripartite structure including a pair of inverted repeats (IRa and IRb) regions of 25,675bp that was separated by a large single copy (LSC) region of 90,255bp, and a small single copy (SSC) region of 18,892bp (Table 1 and Fig.1). The overall GC content of the chloroplast genome was 36.82%, and has a similar GC content to those of other Fagaceae species. Whereas, few differences in the GC contents were found among the chloroplast genome. The contents of the LSC, SSC, and IR regions are 34.65%, 30.94%, and 42.78% respectively (Table 2). The GC content is highest in the IR regions (42.78%), the phenomena was found in the chloroplast genome of *C. hainanensis* [30]. The overall GC content is an important indicator of species affinity[31].

**Table 1.** Summary of four *Castanopsis* chloroplast genome Characteristics.

| Genome | *C. sclerophylla* | *C. hainanensis* | *C. echinocarpa* | *C. concinna* |
| --- | --- | --- | --- | --- |
| Genome size (bp) | 160,497 | 160,631 | 160,647 | 160,606 |
| LSC length (bp) | 90,255 | 90,328 | 90394 | 90,368 |
| SSC length (bp) | 18,892 | 18,929 | 18,995 | 18,884 |
| IR length (bp) | 25,675 | 25,687 | 25,629 | 25,677 |
| Number of genes | 131 | 132 | 132 | 136 |
| Number of protein-coding genes | 86 | 84 | 84 | 82 |
| Number of tRNA genes | 37 | 40 | 40 | 46 |
| Number of rRNA genes | 8 | 8 | 8 | 8 |

**Table 2.** Base content of the *C. sclerophylla* chloroplast genome.

| Region | A (%) | T (%) | C (%) | G (%) | A+T (%) | G+C (%) |
| --- | --- | --- | --- | --- | --- | --- |
| LSC | 31.94 | 33.4 | 17.74 | 16.91 | 65.34 | 34.65 |
| SSC | 34.4 | 34.66 | 16.29 | 14.65 | 69.06 | 30.94 |
| IR | 28.61 | 28.61 | 21.39 | 21.39 | 57.22 | 42.78 |
| Total | 31.65 | 32.23 | 18.47 | 17.65 | 63.18 | 36.82 |

In the *C. sclerophylla* chloroplast genome, a total of 131 genes were found, including 86 protein-coding genes, 37 tRNA genes, and 8 rRNA genes (Fig.1, Table 1). All of these 131 genes, 110 genes were unique and annotated, and were divided into three categories including 79 protein-coding genes, 27 tRNA genes, and four rRNA genes (Table 3). In addition, 21 functional genes (seven protein-coding genes, four rRNA, and 10 tRNA genes) are duplicated in the IR regions (Fig.1). The LSC region comprises 62 protein-coding genes and 22 tRNA genes, whereas the SSC region comprises 11 protein-coding genes and one tRNA gene (Table S1). There are 14 intron-containing genes, including eight protein-coding genes and six tRNA genes. 12 genes contain one intron, with *clpP* and *ycf3* genes having two introns. The *trnK-UUU* gene has the longest intron (2511bp), and the *trnL-UAA* gene has the smallest (485bp) (Table 4). The similar phenomenon is also present in *Quercus acutissima*[32]. *The ycf3* gene has been shown to stable accumulation of photosystem I complexes[33]. So, we will focus on the *ycf3* intron gain of *C. sclerophylla* may be helpful for further study the photosynthesis mechanism.

**Table 3.**
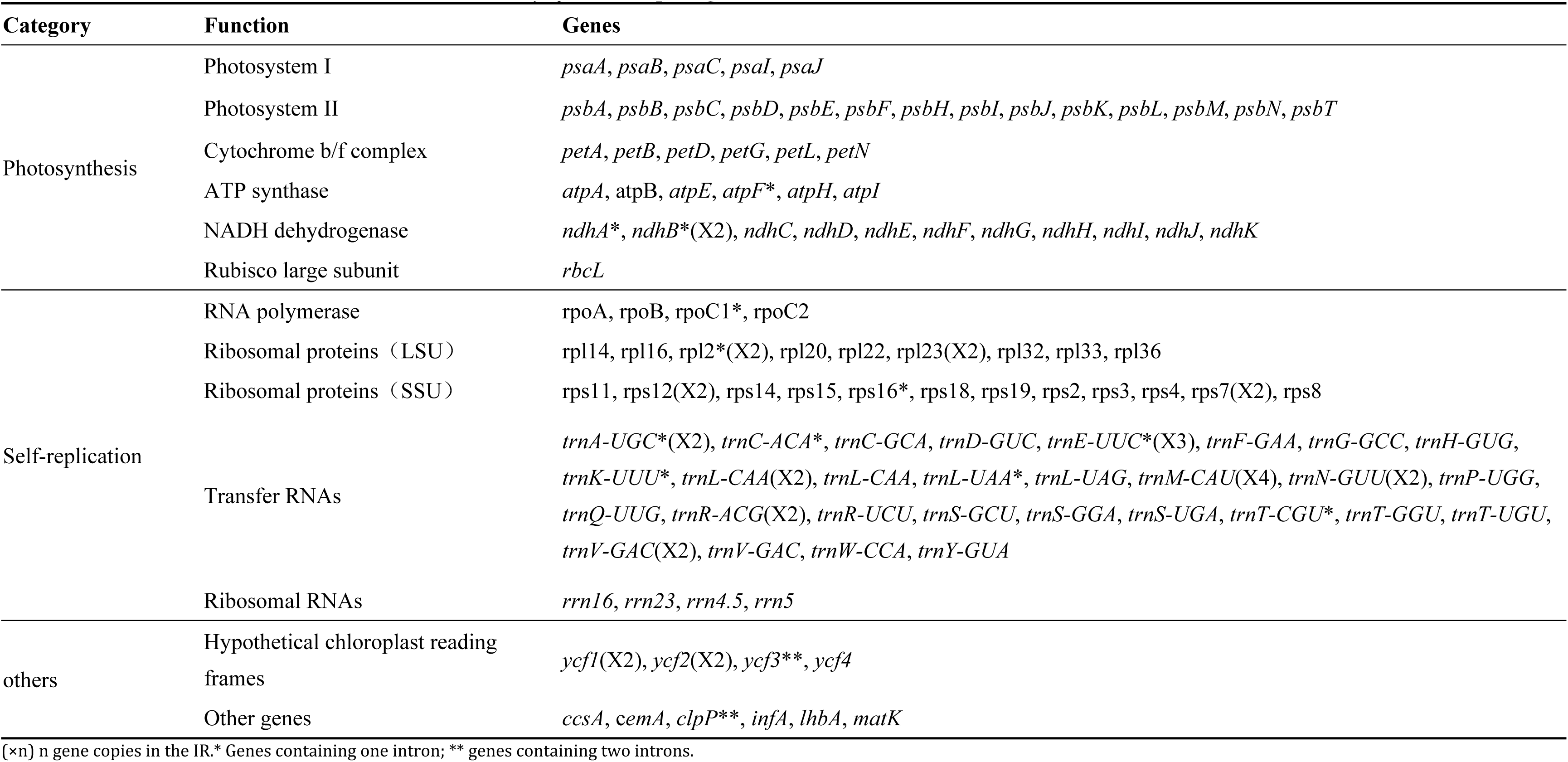
List of genes annotated in the sequenced *C. sclerophylla* chloroplast genome chloroplast genome.

**Table 4.** The lengths of exons and introns in genes with introns in the *C. sclerophylla* chloroplast genome.

(×n) n gene copies in the IR.\* Genes containing one intron; \*\* genes containing two introns.
|  |  |  |  |  |  |  |
| --- | --- | --- | --- | --- | --- | --- |
| Gene | Location | Exon I (bp) | Intron I (bp) | Exon II (bp) | Intron II (bp) | Exon III (bp) |
| *clpP* | LSC | 70 | 851 | 291 | 654 | 227 |
| *trnK-UUU* | LSC | 36 | 2511 | 34 |  |  |
| *rpoC1* | LSC | 429 | 838 | 1618 |  |  |
| *trnC-ACA* | LSC | 37 | 610 | 55 |  |  |
| *ndhB* | IRA | 776 | 681 | 755 |  |  |
| *ndhA* | SSC | 550 | 1049 | 540 |  |  |
| *rpl2* | IRB | 390 | 685 | 433 |  |  |
| *trnA-UGC* | IRB | 36 | 801 | 35 |  |  |
| *trnL-UAA* | LSC | 34 | 485 | 49 |  |  |
| *trnE-UUC* | LSC | 31 | 956 | 39 |  |  |
| *trnT-CGU* | LSC | 34 | 720 | 42 |  |  |
| *rps16* | LSC | 41 | 903 | 227 |  |  |
| *ycf3* | LSC | 125 | 727 | 225 | 768 | 154 |
| *atpF* | LSC | 144 | 789 | 409 |  |  |

**Fig. 1.**
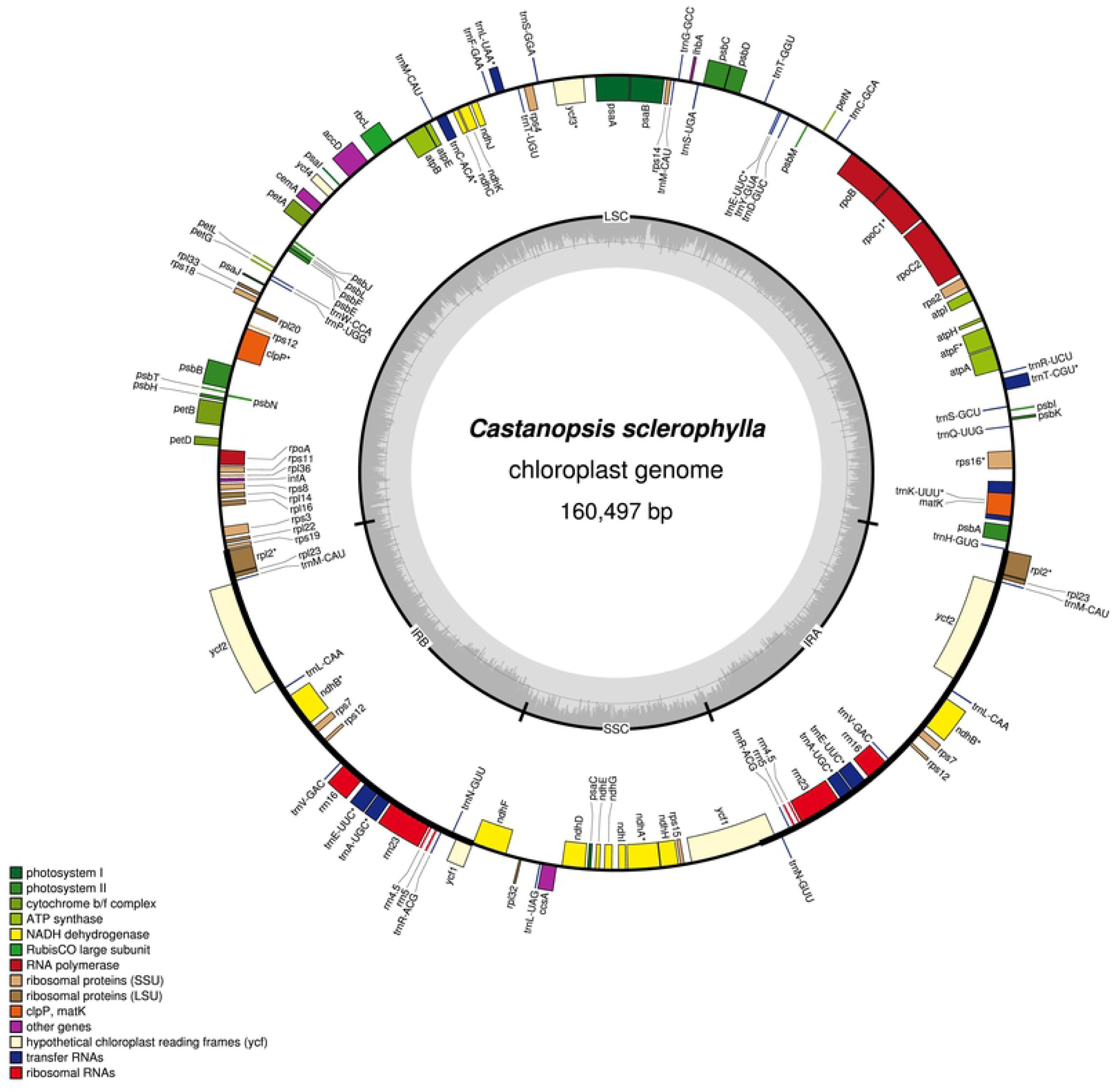
Chloroplast genome annotation map of *C. sclerophylla*. Genes inside the circle are transcribed in a clockwise, whereas genes outside are transcribed in a counterclockwise direction. Different colors represent different functional Genes. The darker gray and lighter gray in the inner circle shows the GC content and AT content of chloroplast genome, respectively.

According to the protein-coding sequences and tRNA genes, the codon usage frequency was computed for *C. sclerophylla* chloroplast genome (Table 5). All genes are encoded by 23131 codons. Among these, leucine (10.61%) is the maximum commonly encoded amino acid, with 2454 codons, and cysteine (1.13%) is the minimum commonly encoded amino acid with 262 codons. Similar ratios for amino acids are found in previously studied chloroplast genomes [34, 35]. The codons ending with A and U are common. Furthermore, the most preferred synonymous codons which relative synonymous codon usage values (RSCU) were bigger than 1 end with A or U[36].

**Table 5.** The codon–anticodon recognition pattern and codon usage for *C. sclerophylla* chloroplast genome.

| Amino acid | Codon | No. | RSCU | tRNA | Amino acid | Codon | No. | RSCU | tRNA |
| --- | --- | --- | --- | --- | --- | --- | --- | --- | --- |
| Ala | GCC | 196 | 0.61 |  | Asn | AAU | 858 | 1.55 |  |
| Ala | GCG | 158 | 0.49 |  | Asn | AAC | 250 | 0.45 |  |
| Ala | GCU | 581 | 1.81 | *trnA-UGC* | Pro | CCG | 136 | 0.56 |  |
| Ala | GCA | 348 | 1.08 |  | Pro | CCA | 270 | 1.12 |  |
| Cys | UGU | 189 | 1.44 | *trnC-ACA* | Pro | CCU | 362 | 1.5 | *trnP-UGG* |
| Cys | UGC | 73 | 0.56 | *trnC-GCA* | Pro | CCC | 195 | 0.81 |  |
| Asp | GAC | 176 | 0.38 |  | Gln | CAG | 179 | 0.44 |  |
| Asp | GAU | 740 | 1.62 |  | Gln | CAA | 638 | 1.56 |  |
| Gln | GAG | 291 | 0.47 |  | Arg | CGA | 312 | 1.38 |  |
| Gln | GAA | 950 | 1.53 |  | Arg | CGC | 94 | 0.41 |  |
| Phe | UUU | 876 | 1.33 | *trnL-UAA* | Arg | CGG | 99 | 0.44 |  |
| Phe | UUC | 438 | 0.67 | *trnN-GUU* | Arg | CGU | 293 | 1.29 | *trnR-ACG* |
| Gly | GGG | 282 | 0.7 |  | Arg | AGA | 418 | 1.84 |  |
| Gly | GGC | 191 | 0.47 | *trnG-GCC* | Arg | AGG | 145 | 0.64 |  |
| Gly | GGU | 526 | 1.3 |  | Ser | UCA | 339 | 1.2 |  |
| Gly | GGA | 615 | 1.52 |  | Ser | UCC | 285 | 1 | *trnS-GGA* |
| His | CAU | 419 | 1.54 |  | Ser | AGC | 104 | 0.37 |  |
| His | CAC | 126 | 0.46 |  | Ser | UCU | 474 | 1.67 | *trnT-UGU* |
| Ile | AUC | 384 | 0.56 |  | Ser | AGU | 345 | 1.22 |  |
| Ile | AUA | 664 | 0.97 |  | Ser | UCG | 155 | 0.55 | *trnT-CGU* |
| Ile | AUU | 1000 | 1.46 |  | Thr | ACU | 474 | 1.61 |  |
| Lys | AAA | 920 | 1.5 |  | Thr | ACC | 217 | 0.74 |  |
| Lys | AAG | 306 | 0.5 |  | Thr | ACA | 355 | 1.2 |  |
| Leu | CUG | 170 | 0.42 |  | Thr | ACG | 133 | 0.45 |  |
| Leu | UUA | 810 | 1.98 |  | Val | GUG | 180 | 0.56 |  |
| Leu | CUA | 316 | 0.77 |  | Val | GUU | 461 | 1.43 | *trnE-UUC* |
| Leu | CUU | 496 | 1.21 | *trnL-UAG* | Val | GUC | 154 | 0.48 | *trnV-GAC* |
| Leu | CUC | 170 | 0.42 | *trnH-GUG* | Val | GUA | 499 | 1.54 |  |
| Leu | UUG | 492 | 1.2 | *trnM-CAU* | Trp | UGG | 402 | 1 | *trnW-CCA* |
| Met | AUG | 533 | 1 |  | Tyr | UAC | 179 | 0.41 |  |
|  |  |  |  |  | Tyr | UAU | 690 | 1.59 |  |

Note: RSCU indicated Relative Synonymous Codon Usage.

### Comparative Analysis of Genomic Structure

Three complete chloroplast genomes within the *Castanopsis* genus (*C. hainanensis, C. echinocarpa*, and *C. concinna*) were selected for comparison with that of *C. sclerophylla*. *C. sclerophylla* has the smallest chloroplast genome (160,497bp), whereas *C. echinocarpa* has the largest chloroplast genome (160,647) with the smallest IR region (25,629 bp). Additionally, the LSC region lengths varied in these four species, from 90,255 bp of *C. sclerophylla* to 90,394 bp of *C. echinocarpa* (Table 1). The different lengths of LSC region are the main reason for four specie in sequence lengths, this result is coincident with the genus of *Oryza*[34]. *To investigate the levels of the genome divergence, sequence identity was plotted for four species chloroplast genomes with C. concinna* as a reference by the program mVISTA (Fig.2). The results of sequence analysis detected that the LSC and SSC regions were more divergent than the IR regions among the four genomes within the *Castanopsis*, furthermore, a higher divergence was found in non-coding regions than in coding regions. Significant variation in coding regions of the four chloroplast genomes were *ndhF, ndhG*, and *ycf1*, which were all located in the SSC regions. However, the most divergent regions were observed in the intergenic regions, including *trnK*-*rps16, trnS*-*trnT, atpA*-*atpF, trnC*-*petN, trnT*-*psbD, IhbA*-*trnG, ycf3*-*trnS, rps4*-*trnT, trnT*-*trnL, atpB*-*rbcL, petA*-*psbJ, psbE*-*petL, rpl16*-*rpl3, ndhF*-*rpl32*.

The expansion and contraction of IR regions at the borders are the major reason for chloroplast genome size variations and play vital roles in evolution [37–39]. A detailed comparison of four junctions (JLA, JSB, JSA, and JLA) between the two single-copy regions (LSC and SSC) and the two IRs (IRa and IRb) was performed among *C. sclerophylla, C. hainanensis, C. echinocarpa* and *C. concinna* by analyzing the exact IR border positions and adjacent genes (Fig.3). The IR regions are relatively conserved in the genus of *Castanopsis*, this result is agreed with the reports in the genus of *Quercus*[32]. The *rpsl9* gene was located between the junction of LSC and IRb regions in *C. concinna*. However, in *C. sclerophylla, C. hainanensis,* and *C. echinocarpa* chloroplast genomes, the *rps19* gene was completely located in LSC region and had an 11bp, 11bp, and 10bp distance to the border of LSC, respectively. Some studies indicate that *ycf1* is requisite for plant viability and encodes Tic214 which is an vital component of TIC complex in *Arabidopsis*[40, 41]. The *ycf1* gene crossed the SSC/IRb and SSC/IRa regions. The SSC/IRb junction is located in the *ycf1* region in all four *Castanopsis* species chloroplast genomes and extends into the SSC region by different lengths depending on the genome (*C. sclerophylla*, 21bp; *C. hainanensi****s***, 24 bp; *C. echinocarpa,* 59 bp; *C. concinna*, 59 bp); the IRb region includes 1131, 1157, 1107, and 1092bp of the *ycf1* gene. The SSC/IRa junction extends into the SSC region by different lengths depending on the genome (*C. sclerophylla*, 4581bp; *C. hainanensis*, 4568 bp; *C. echinocarpa,* 4608 bp; *C. concinna*, 4581bp); the IRa region includes 1092, 1114, 1107, and 1092bp of the *ycf1* gene.

**Fig. 2.**
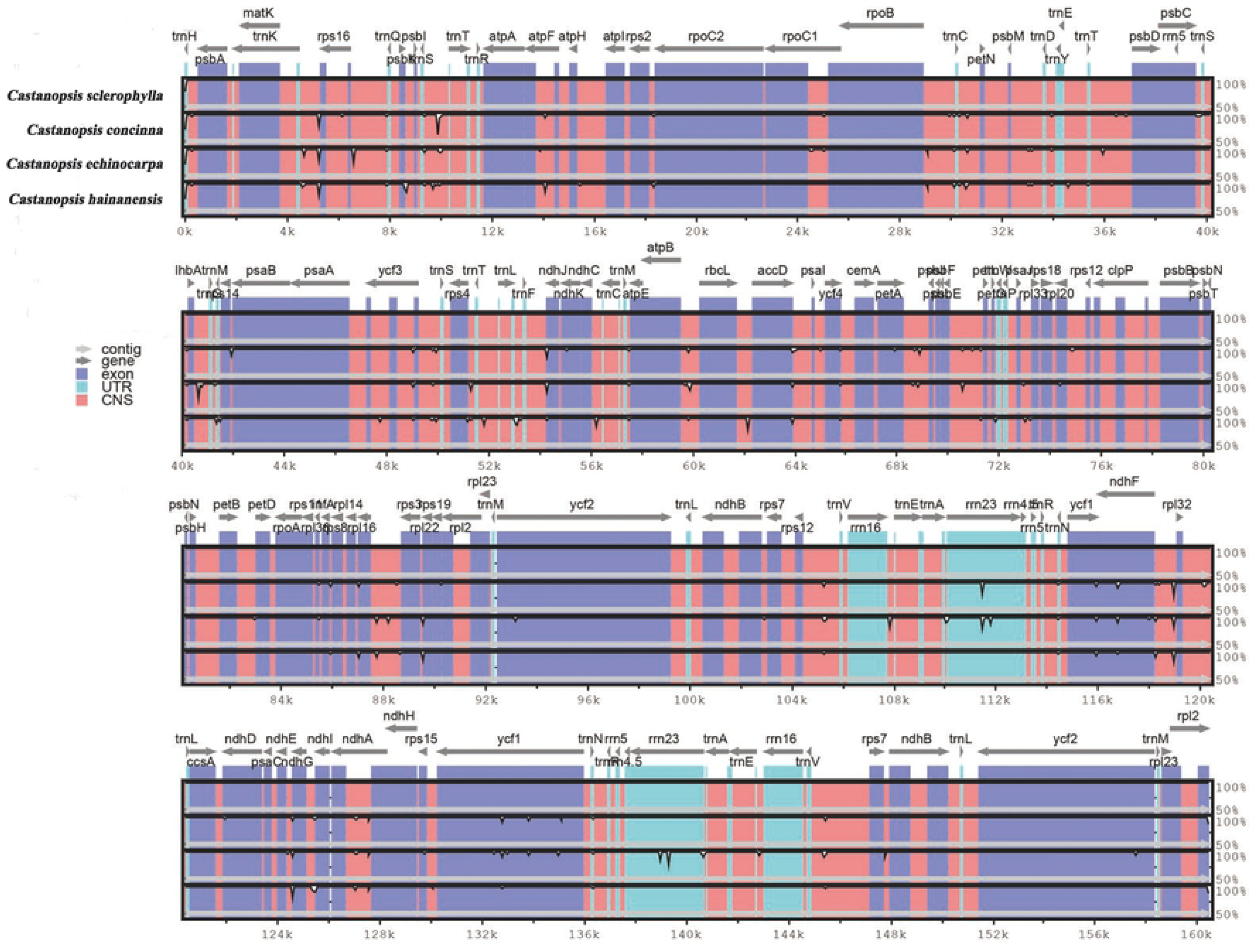
Visualization alignment of the complete chloroplast genome of four species with C. concinna as a reference by the program mVISTA. The grey arrows and thick black lines above the alignment indicate the orientation of genes. Blue bars represent exons, sky-blue ones represent untranslated region (UTR), and pink ones represent non-coding sequences (CNS). The vertical scale represents the percent identity within 50–100%.

**Fig. 3.**
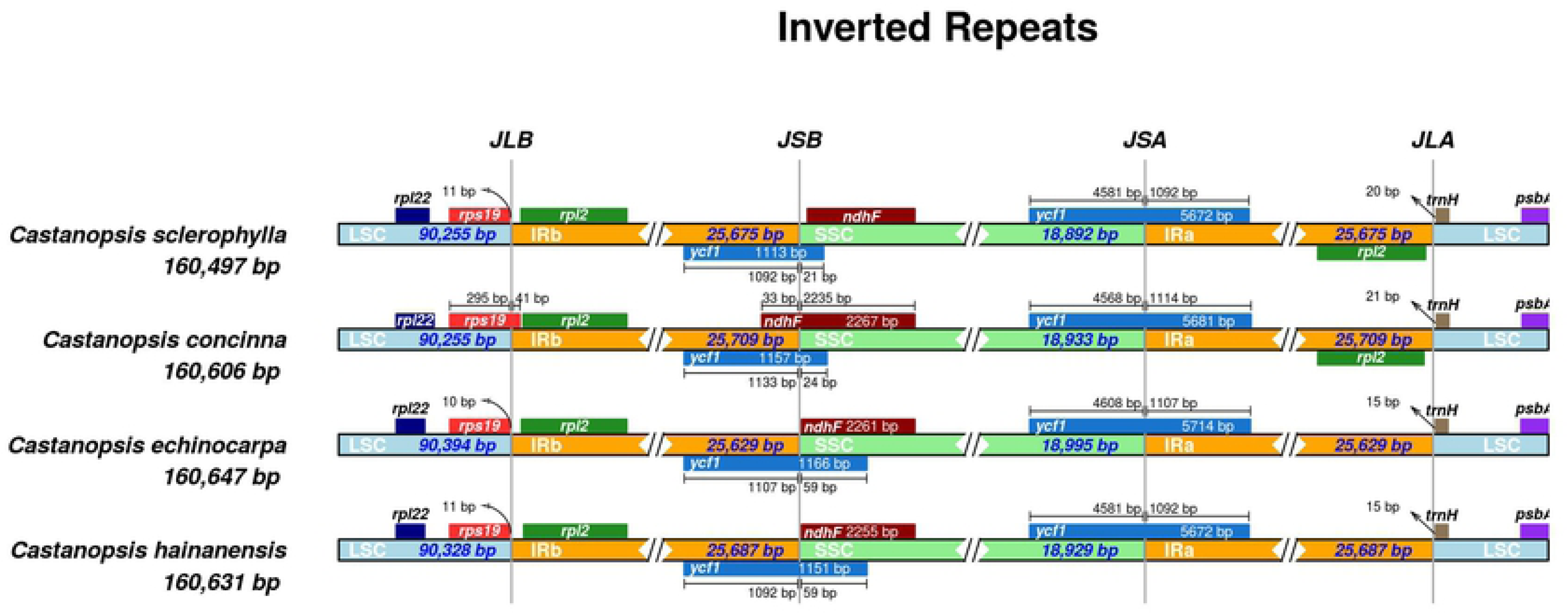
Comparison of junctions the large single copy (LSC), small single copy (SSC), and inverted repeat (IR) among four chloroplast genomes of congeneric species.

### Phylogenetic Analysis

Phylogenetic analysis was constructed based on the 22 alignment sequences chloroplast genomes by maximum likelihood (ML) analysis (Fig. 4). *Corylus fargesii* and *Eucalyptus umbra* were used as the outgroups. The maximum likelihood (ML) phylogenetic analysis showed that these four species of the genus *Castanopsis* formed a monophyletic clade and that *C. sclerophylla* is closely related to *Castanopsis hainanensis* with strong bootstrap values. The ML tree indicated that *Castanopsis* was closely related to *Castanea*. Surprisingly, the species of *Quercus* genus were not formed a clade, in addition, this genus *Quercus* were not divided to two clusters according to evergreen tree species deciduous tree species. The phylogenetic status of these genus were consistent with the previous report[32]. Until now, little has been known about the chloroplast genome of the *Castanopsis*, and only three chloroplast genome sequences of the *Castanopsis* species are found in GeneBank, which has greatly hampered for studying the phylogenetic relationships of *Castanopsis*. So, it is necessary for more research of the complete chloroplast genome within the *Castanopsis* genus in the future.

**Fig. 4.**
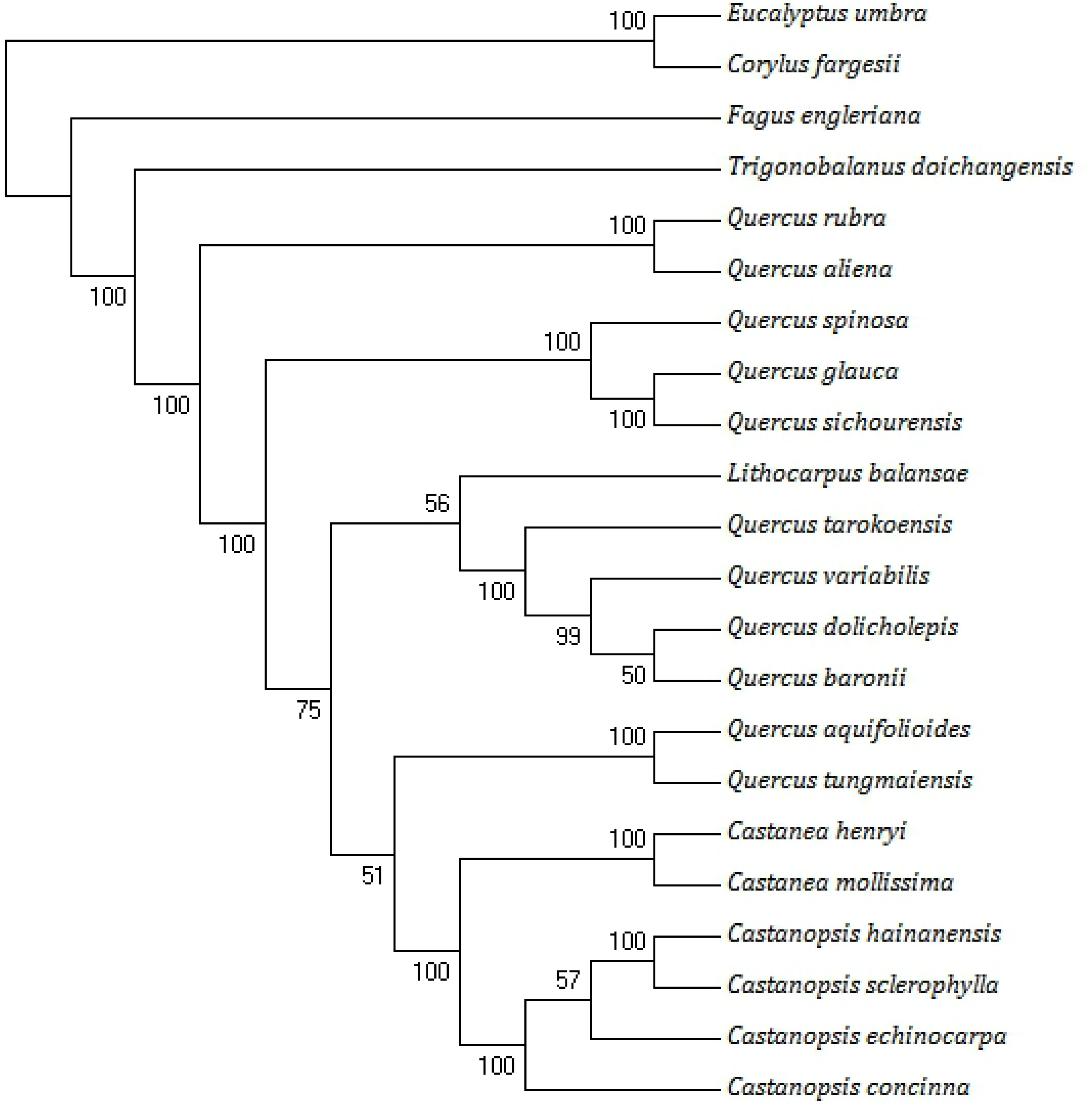
A maximum likelihood (ML) phylogenetic tree was constructed based on 22 species chloroplast genomes. *Corylus fargesii* and *Eucalyptus umbra* were used as the outgroups.

## Conclusions

*C. sclerophylla* is an important evergreen broad-leaved species of the *Castanopsis* genus in the Fagaceae family. In this study, the complete chloroplast genome sequences of C. *Castanopsis* sclerophylla was reported based on the Illumina Hiseq 2500 platform. *C. sclerophylla* chloroplast genomes exhibited a typical quadripartite and circular structure, which is similar to other three congeneric species. Compared to the chloroplast genomes of three *Castanopsis* species, *C. sclerophylla* has the smallest chloroplast genome (160,497bp). The phylogenetic ML tree showed that Phylogenetic relationships among 22 angiosperms strongly supported the known classification of *C. sclerophylla*. The ML analysis showed that these four species of the genus *Castanopsis* formed a monophyletic clade and that *C. sclerophylla* is closely related to *C. hainanensis* with strong bootstrap values, in addition, *Castanopsis* was closely related to *Castanea*. The genus *Castanopsis* contains about 120 known species, nearly half of these species are native to China. China has plenty of *Castanopsis* germplasm resources, the availability of the chloroplast genomes provides a powerful genetic resource for the phylogenetic analysis and biological study. So, it is necessary for research of the complete chloroplast genome of the genus *Castanopsis* in the future. The data will contribute to understand the genetic resources and identification of evolutionary relationships, and facilitate to the exploration, utilization and conservation genetics of the genus.

## Acknowledgments

We are grateful to Yuelong Liang for his assistance in sample collection.

## Supporting information

**S1 Table.** The number of genes *C. sclerophylla* chloroplast genome chloroplast genome

| Region | Number of CDS | Number of tRNA | Number of rRNA | Total |
| --- | --- | --- | --- | --- |
| LSC region | 62 | 22 | 0 | 84 |
| SSC region | 11 | 1 | 0 | 12 |
| IRA region | 7 | 7 | 4 | 18 |
| IRB region | 6 | 7 | 4 | 17 |

